# Novelty and uncertainty interact to regulate the balance between exploration and exploitation in the human brain

**DOI:** 10.1101/2021.10.13.464279

**Authors:** Jeffrey Cockburn, Vincent Man, William Cunningham, John P. O’Doherty

## Abstract

Recent evidence suggests that both novelty and uncertainty act as potent features guiding exploration. However, these variables are often conflated with each other experimentally, and an understanding of how these attributes interact to regulate the balance between exploration and exploitation has proved elusive. Using a novel task designed to decouple stimulus novelty and estimation uncertainty, we identify separable behavioral and neural mechanisms by which exploration is colored. We show that uncertainty was avoided except when the information gained through exploration could be reliably exploited in the future. In contrast, and contrary to existing theory, novel options grew increasingly attractive relative to familiar counterparts irrespective of the opportunity to leverage their consequences and despite the uncertainty inherent to novel options. These findings led us to develop a formal computational framework in which uncertainty directed choice adapts to the prospective utility of exploration, while novel stimuli persistently draw favor as a result of inflated reward expectations biasing an exploitative strategy. Crucially, novelty is proposed to actively modulate uncertainty processing, effectively blunting the influence of uncertainty in shaping the subjective utility ascribed to novel stimuli. Both behavioral data and fMRI activity sampled from the ventromedial prefrontal cortex, frontopolar cortex and ventral striatum validate this model, thereby establishing a computational account that can not only explain behavior but also shed light on the functional contribution of these key brain regions to the exploration/exploitation trade-off. Our results point to multiple strategies and neural substrates charged with balancing the explore/exploit dilemma, with each targeting distinct aspects of the decision problem to foster a manageable decomposition of an otherwise intractable task.

## 1 Introduction

Adaptive organisms are faced with a fundamental trade-off between choosing a familiar option that leads to a known reward, or exploring less familiar alternatives in hopes of finding something better [1]. The explore-exploit dilemma presents an exceptional challenge with only a narrow range of circumstances in which an optimal solution is known [2]. Yet, despite its importance to survival, very little is known about how the human brain tackles this situation, or how those neural computations manifest themselves behaviorally. However, an emerging literature has highlighted the contributions of two variables driving exploration in the mammalian brain: stimulus novelty [3, 4, 5, 6], and outcome uncertainty [7, 8, 9, 10, 11]. Despite their significance, these variables are often conflated, and nothing is known about how they interact to regulate exploration. Here, we describe a bespoke behavioral task specifically designed to distinguish these two variables from each other, thereby allowing both their unique and interactive influence on behavior to be assessed. We craft and empirically test a new computational framework to describe precisely how these variables contribute to exploration using behavioral and neural data measured using functional magnetic resonance imaging (fMRI).

The machine learning literature offers a growing catalogue of practical approaches to the explore/exploit dilemma. Based on the assumption that uncertainty often points to learning opportunities, one class of algorithms tries to improve sampling efficiency by directing the decision making process according to an uncertainty bonus [12, 13, 14]. An alternative strategy, commonly referred to as optimistic value initiation, boosts the initial reward expectation associated with novel options, compelling the learning agent to probe novel opportunities that would not otherwise be favored over more familiar alternatives [15, 16]. As an intriguing point of contrast, algorithms that employ an exploration bonus during each decision can flexibly accommodate motivational changes and environmental volatility, whereas optimistic initiation fuses the exploratory motivation into the reward expectation to foster exploration through a singular exploitative mechanism. We set out to investigate this trade-off between exploratory flexibility and computational efficiency as an additional understudied dimension of human learning and decision making [17].

Studies have investigated human strategies for regulating the explore/exploit trade-off with mixed results. Humans robustly avoid uncertainty when it cannot be reduced or taken advantage of [18], suggesting that uncertainty itself is undesirable. However, varied findings have emerged from learning tasks in which sampling uncertain options can provide beneficial information, with reports of uncertainty aversion [19], indifference [20], and uncertainty seeking behavior [11, 10] amongst a range of individual differences [8, 21]. Notably, when expected value and uncertainty are carefully decoupled, human participants preferentially sample uncertain options when given the opportunity to leverage what they’ve learned [7, 9], suggesting that uncertainty-directed exploration is motivated in part by the prospect of making more rewarding choices in the future. In contrast, several lines of evidence show that animals and humans alike exhibit a robust preference for novel options [3, 4, 5], so much so that that marketing strategies depend on it [22]. A puzzling incongruity is the fact is that while appetites for uncertainty vary across individuals and tasks, novelty robustly draws favor despite the fact that novel options are themselves inherently uncertain.

Research probing the neural correlates of exploration offer additional constraints on how the human brain balances this trade-off. Several studies have implicated frontopolar cortex (FPC) as tracking the relative uncertainty of the options being considered [8, 23], or the potential advantage of switching to an alternative course of action [24, 25], and is uniquely engaged when exploratory choices are made [20]. Disrupting FPC using transcranial magnetic stimulation (TMS) increases (stimulation) or decreases (inhibition) uncertainty directed choice [26], further implicating its role in exploration. Additional studies have also highlighted the ventral medial prefrontal cortex (vmPFC), a region associated with both value guided choice [27] and outcome monitoring[28], as mitigating the switch between exploratory and exploitative phases of behavior [10, 11, 29], suggestive of a multi-hub circuit concerned with balancing exploration and exploitation.

Studies investigating the neural underpinnings of novelty processing have shown that dopamine (DA) neurons elicit a phasic burst when novel stimuli are experienced [30]. Consistent with a rich line of research linking DA to the reward prediction error learning signal (RPE), where unexpected reward or stimuli predictive of reward induce phasic DA signals [31, 32], it has been suggested that the novelty induced phasic burst of DA reflects a shaping bonus encouraging exploration [33]. Imaging results support this idea, showing that the ventral striatum reflects a skewed RPE signals consistent with optimistic initiation [6, 34].

Although it has been established that the factors driving exploration are multi-faceted, little is known about how these features coexist and interact, and in particular, how the tension between novelty and uncertainty is resolved. Complicating this issue enormously is the fact that novelty and uncertainty are challenging to distinguish experimentally because novel options tend to be those that are maximally uncertain, and a natural correlation emerges as both novelty and uncertainty decline with additional engagement. However, we hypothesized that it would be possible to uncover the unique influence of these variables by decoupling their respective contributions to learning and decision making as a function of task horizon. Specifically, we hypothesized that novelty operates as a simple exploratory heuristic that guides approach behavior via optimistic initiation, and since this mechanism colors exploration by way of reinforcement learning, the effects of novelty should persist without considering the task’s horizon. On the other hand, the more computationally rich variable of estimation uncertainty is hypothesized to reflect the prospective value of information gain, and as such, should adaptively accommodate task horizon to render uncertain options less desirable with diminishing opportunity to leverage what might be learned about them.

To test these ideas, we exposed human participants to a newly designed bespoke decision making task while undergoing fMRI. The task consisted of a series of games in which participants were asked to choose between options that varied in terms of novelty, uncertainty and expected reward. Our task design offers two important experimental advances. First, it dissociates uncertainty and novelty by explicitly revaluing options, thereby increasing uncertainty without affecting novelty. Second, novel and uncertain options were systematically offered as a function of proximity to a game’s termination, allowing us to probe feature specific adaptation as a function of task horizon.

We develop a comprehensive computational framework to describe how novelty and uncertainty interact and guide behavior. Importantly, given the subtle behavioral distinctions that emerge from various model implementations, we leverage fMRI data to validate and discriminate between different algorithmic forms of the model by focusing on distinguishable patterns of activity in key regions of interest identified based on their role in implementing exploration/exploitation computations, namely the ventral striatum, the ventromedial prefrontal cortex and the frontal pole.

## 2 Results

Participants (n=32) performed a novel learning and decision making task while undergoing fMRI. In brief, participants played 20 blocks of a finite horizon multi-armed bandit task designed to expose the influence of reward history, estimation uncertainty and stimulus novelty on balancing the explore/exploit trade-off while undergoing four consecutive 15-minute fMRI sessions (5 learning contexts blocks per session). On each trial, participants were asked to choose between two slot machines. Having made their choice they were informed of the machine’s payout of either $1US (win) or $0US (loss). Participants were instructed that each learning context block, framed as a visit to a new casino, would consist of approximately 20 trials (between 18 to 23 trials) and that they should try to accumulate as much reward as they could. At the end of the experiment, one of the casinos they visited was chosen at random from which a performance bonus was calculated.

Each block offered a new learning context that included five visually unique and identifiable slot machines, three of which were familiar stimuli that had been seen in previously visited casinos, and the remaining two were novel stimuli that had not yet been shown (43 stimuli in total). Participants were informed that each slot machine had a fixed probability of winning within a given casino, but all machines were programmed differently by each casino, and as such, anything learned in one casino should not be applied to others. Trials were structured such that the two slot machines varied in terms of reward probability (expected value manipulation), the number of previous exposures (novelty manipulation), as well as the number of times they’d been sampled in the current block (uncertainty manipulation), allowing us to to systematically examine the influence of reward, novelty and uncertainty across the task horizon.

### 2.1 Choice reflects reward history, novelty, and uncertainty

We first examine the degree to which reward history, uncertainty, and stimulus novelty influence choice. To investigate the influence of reward history, choice was modelled as a function of the difference in expected value between the left and right options (*EV*_*L*_− *EV*_*R*_), where expected value was defined as the mean of a Beta distribution specified according to each option’s history of wins and losses (Beta(*α* =number of wins + 1, *β* = number of losses + 1)). As illustrated in Figure 2A, choice was robustly governed by reward history, as exhibited by an indifference between equally rewarded options (*β*_*EV* 1=*EV* 2_ = 0.02, *p* = 0.73), and an increasingly reliable preference for the option with a higher expected value (*β*_*EV*1−*EV*2_ = 4.76, *p* < 0.01). Importantly, value learned in previous contexts did not influence behavior, demonstrating that participants were motivated and understood the task structure (see Supplement for details).

**Figure 1:**
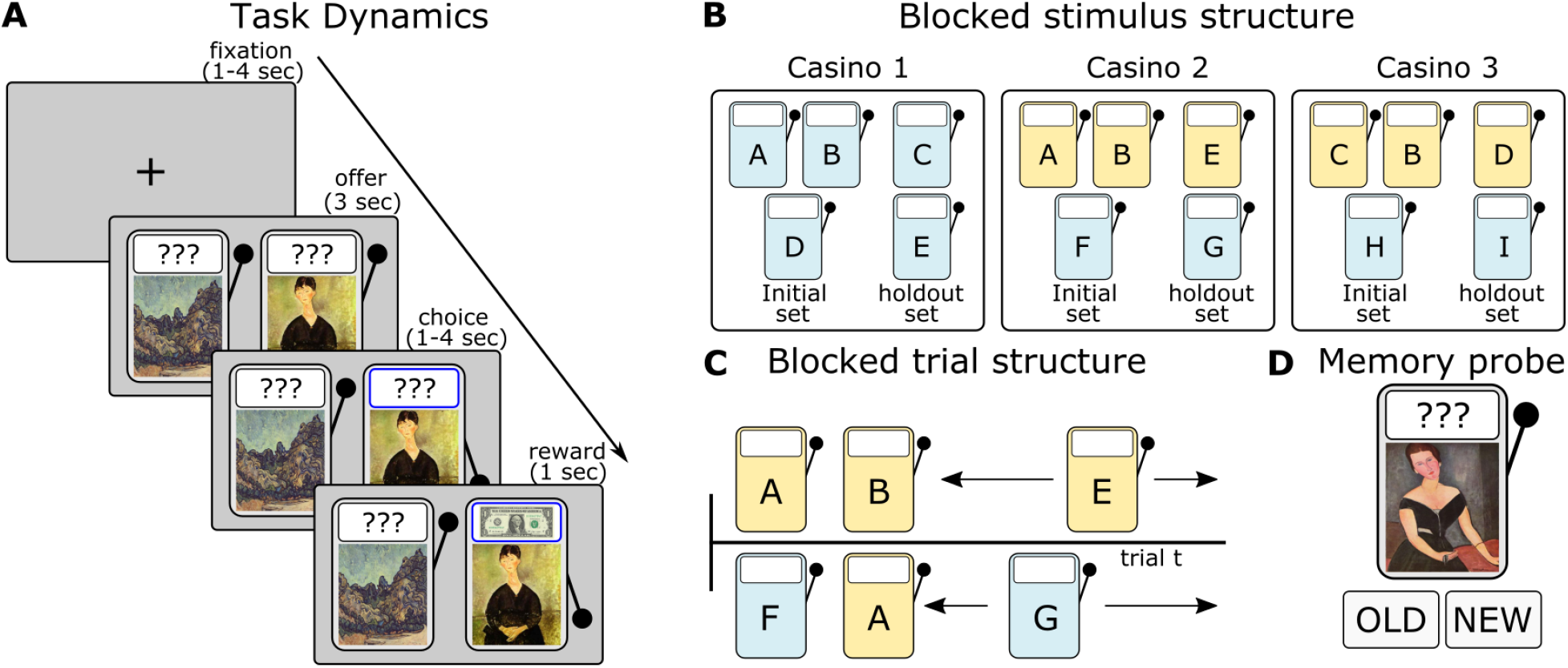
Experimental procedures. **A)**: Trial dynamics. Following a jittered fixation screen (sampled from a pseudo-randomized linear spacing of 1-4 seconds), participants were offered two slot machines, and asked to select one using a button box (left thumb to select the option on the left, right thumb to select the option on the right). Choice feedback was provided (handle moved to a pulled position, and outcome frame highlighted). Following a 1-4 second jitter (sampled from a pseudo-randomized linear spacing of 1-4 seconds), reward feedback ($1/$0) was shown. **B)** Blocked stimulus structure: Each learning context was framed as a visit to a unique casino. Five different slot machines could be offered in each casino, two of which were novel and three were familiar (coloured blue and yellow respectively here for demonstration purposes - all machines using during the task were grey). Stimuli were segregated to form an initial set of three stimuli used early in each block, and a holdout set of two stimuli (one novel, the other familiar) that were gradually added to the initial set from which offered stimuli may be drawn on each trial. **C)** Blocked trial structure: Using stimuli from the initial set, pairs of offers were made. This included pairs with equal uncertainty but differing familiarity, as well as pairs with equal familiarity but differing uncertainty. At pseudo random trials, the novel and familiar holdout stimuli were added to the set from which stimuli may be drawn. **D)** Memory probe task: Having completed 20 blocks of the casino task, participants were asked to label stimuli as old (they had seen them in the casino task) or new.

**Figure 2:**
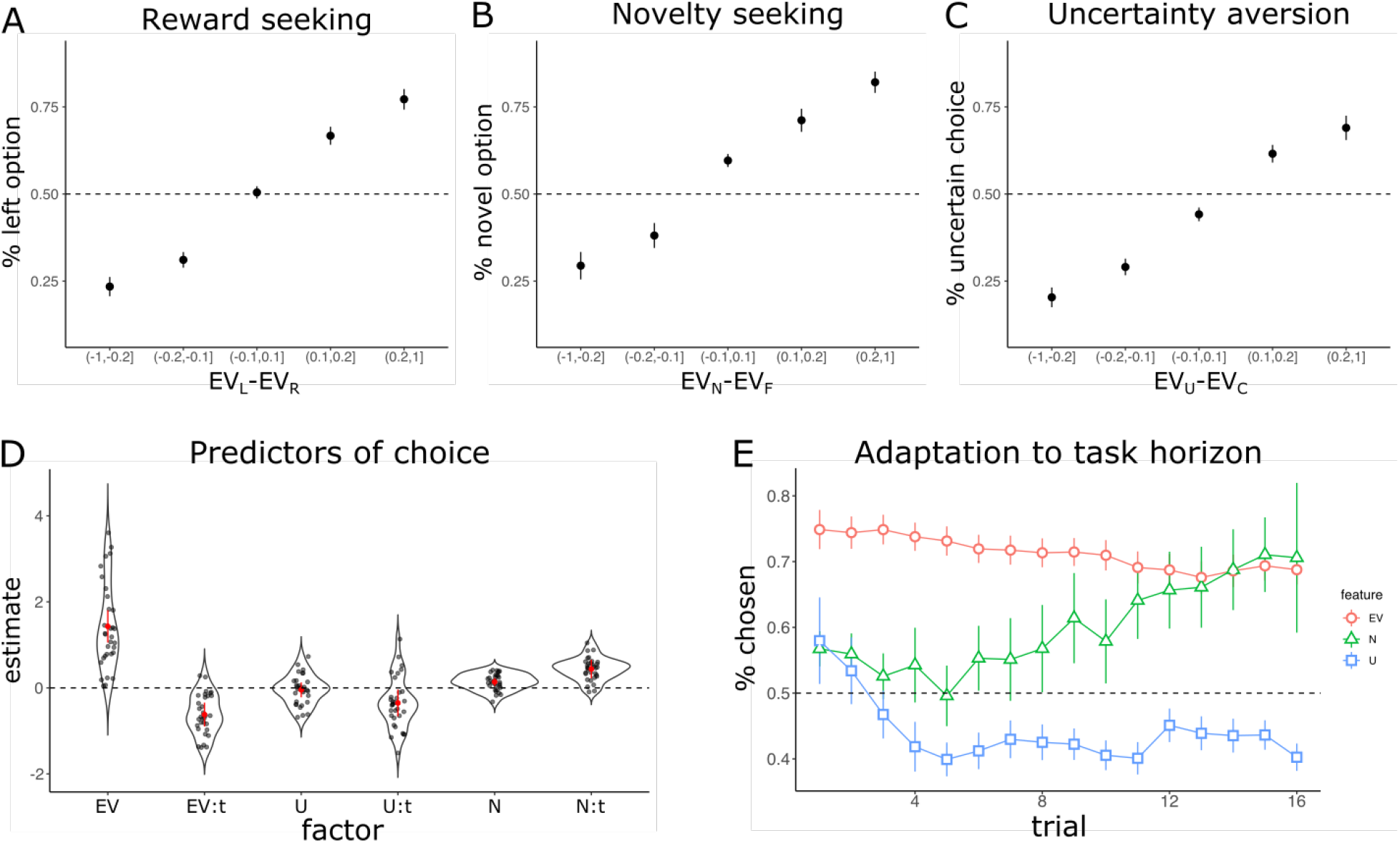
Behavioral evidence for the influence of reward history, novelty and uncertainty on choice. **A)** The proportion of trials in which the left option was chosen as a function of the difference in expected value between the two options (*EV*_*L*_ − *EV*_*R*_). **B)** The proportion of trials in which the novel option was chosen as a function of the difference in expected value between the novel and familiar option (*EV*_*N*_ − *EV*_*F*_) **C)** The proportion of trials in which the uncertain option was chosen as a function of the difference in expected value between the uncertain and more certain option (*EV*_*U*_ − *EV*_*C*_) **D)** Estimates from a logistic regression showing the influence of stimulus features on choice, and their trajectory across a block of trials. Points depict random effects, red points and error bars depict fixed effects and 95% confidence intervals respectively **E)** The proportion of trials in which the option with higher expected value was selected (red), the option that was most novel was selected (green), and the option that was most uncertain was selected (blue) across the task horizon. Points in panels A-C represent mean scores across participants, and error bars depict standard error of the participant mean scores

Next, we probed the influence of stimulus novelty by focusing on the subset of trials in which a novel option was offered, where a stimulus is considered novel if it had been presented fewer than three times prior to the current trial. The proportion of trials in which the novel option was sampled as a function of the alternative option’s expected value is illustrated in Figure 2B. While reward history exerts the dominant force 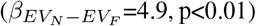, participants exhibited a robust novelty-seeking bias when both options were of approximately equal value 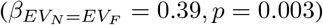.

We revisited this analysis to investigate the effect of uncertainty, which we quantify as the variance of the Beta distribution describing stimulus outcome history. Limiting our analysis to the subset of trials in which participants were offered familiar options with unequal variance (Figure 2C), again, we see that choice is strongly shaped by reward history 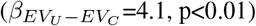. However, in contrast to the observed effects of novelty, participants shied away from more uncertain option when the expected value was approximately equal 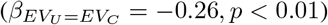.

### 2.2 The influence of reward history, stimulus novelty and estimation uncertainty adapted to task horizon

Our experimental design allows us to probe how reward history, novelty and uncertainty differentially influence choice across the learning context horizon. Here, we characterized these effects using a computationally agnostic logistic regression analysis (glmer in lme4) to model choice on each trial. Again, we defined expected value as the mean of a Beta distribution specified according to each option’s history of wins and loses in the current context. Uncertainty was defined as the variance of the same distribution, and stimulus novelty was defined as the variance of a Beta distribution specified according to the total number of exposures (Beta(*α* = number exposure + 1, *β* = 1)). Choice on each trial was then modelled as a function of the difference in expected value (*EV*_*L*_ − *EV*_*R*_), stimulus novelty (*N*_*L*_ − *N*_*R*_), and uncertainty (*U*_*L*_ − *U*_*R*_). Lastly, we probed for feature specific adaptation across the horizon of a learning context by including trial number as an interaction term with each feature of interest.

As illustrated in Figure 2D, reward history is indeed a strong predictor of choice (*β*_*EV*_ = 1.43, *p* < 0.01), with participants preferring the option with a higher expected value. However, the influence of expected value diminished as participants progressed through a block of trials (*β*_*EV*:*t*_ = −0.62, *p* < 0.01), suggesting some deviation from optimal outcome integration and value exploitation. Uncertainty also had a significant effect on choice. Participants expressed differing strategies at the start of a learning block, with a roughly even split of participants directing their sampling towards more uncertain options or trying to avoiding them, resulting in a group average that did not differ from zero (*β*_*U*_ = −0.04, *p* < 0.63). However, a model that included uncertainty offered a significant improvement over a model that did not (*χ*^2^(23, 32) = 210, *p* < 0.01), showing that uncertainty played a significant but varied role across individuals early in learning. In line with optimal theories of exploration [2], participants grew increasingly reluctant to explore uncertain options as the probability of task termination increased (*β*_*U*:*t*_ = −0.37, *p* = 0.02). Lastly, participants expressed a robust novelty seeking bias at the start of the learning context (*β*_*N*_ = 0.14, *p* < 0.01). In contrast to the growing uncertainty aversion, novel options grew increasingly attractive as the block of trials unfolded (*β*_*N*:*t*_ = 0.44, *p* < 0.01).

The feature specific trajectories exposed by this analysis are depicted in Figure 2E, showing the proportion of trials in which the stimulus with greater expected value (red), uncertainty (blue), or novelty (green) was chosen as a function of progress through a learning context. Decisions between options with differing expected reward strongly favored that with higher value, though this effect waned as the task proceeded. Trials in which both options were familiar but differed in terms of uncertainty saw participants grow less likely to probe the most uncertain stimulus offered, consistent with the normative exploration strategy in a bandit task such as ours. In contrast, participants expressed an increasing preference for novel options, despite their inherent uncertainty.

### 2.3 A replication of novelty seeking and uncertainty aversion

We sought to establish the reliability of key behavioral signatures highlighted in our fMRI study. Behavioral data was collected from N=79 participants at the University of Toronto, where participants were exposed to a variant of the experimental protocol in which the first 6 blocks of the experiment replicate the task design described here (see methods for details). We report analyses from those 6 initial trial blocks here (the remainder of the experiment focused on affective manipulations which will be reported on elsewhere).

Replicating our original findings, participants in this second study were influenced by reward history, stimulus novelty, and estimation uncertainty. Mirroring the analyses reported in Figure 2A-C, choice was strongly govern by reward history, with participants exhibiting an indifference between equally valued stimuli (*β*_*EV*1=*EV* 2_ =− 0.04, *p*= 0.4), with an increasingly reliable preference for the option with a higher expected value (*β*_*EV*1−*EV*2_ = 3.77, *p* < 0.01). Participants also elicited a strong novelty seeking bias when both options were of approximately equal value 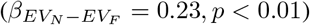, as well as an aversion to more uncertain options 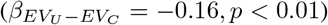.

Noting that participants in the replication study were both novelty seeking and uncertainty averse, we probed the trajectory of each feature’s influence on choice by administering the same regression analysis reported in Figure 2D. We observed similar results to those of the original fMRI study, with participants expressing a growing preference for novel options (*β*_*N*:*t*_ = 0.14, *p* < 0.01), while simultaneously developing an increasing distaste for more uncertain options (*β*_*U*:*t*_ = −0.11, *p* = 0.04). Thus, participants in the replication study also exhibit the same growing tension between novelty and uncertainty.

In summary, using a computationally agnostic regression model of choice we found evidence of both exploration and exploitation during our task, which was replicated in a larger behavioral study. Of note, we identify clear conflict between features associated with exploration and their trajectories across the task at hand, expressed as a growing distaste for uncertainty concomitant with a growing appetite for novel alternatives despite their inherent uncertainty. Using these results to benchmark and constrain our consideration of the mechanisms driving choice we turn to computational models of learning and decision making to elucidate how these patterns emerge, focusing not only on how the trade-off between exploration and exploitation is regulated, but also the puzzling tension between novelty and uncertainty guiding exploration.

### 2.4 Computational model of choice

Regression analyses identified similar choice patterns in both fMRI and replication behavioral data sets, showing that reward history, stimulus novelty, estimation uncertainty, and task horizon all influence choice. Here, we apply computational models of learning and decision making to behavior collected during the fMRI study as a path towards a better understanding of how the explore/exploit trade-off is regulated in the human brain.

The task was modelled from the perspective of a forgetful Bayesian learner. In brief, each option is represented as a Beta distribution, which was defined according to a recency weighted integration of previously observed outcomes. Novelty was accommodated by way of optimistic value initiation by either inflating the initial *α* (novelty seeking) or *β* (novelty averse) parameters used to specify each option’s Beta distribution. From this representation we employ the distribution’s mean and variance as the option’s expected value and uncertainty respectively. In line with theoretical and empirical results, uncertainty was incorporated into the decision making process as a bonus term (see materials and methods for details).

Uncertainty and novelty were both shown to influence choice, but each followed separable trajectories across the task’s horizon (see Figure 2D-E). We accommodated this adaptive influence into the model by including feature weights that consider progress through the learning context. Using free parameters that define the initial (*U*_*I*_, *N*_*I*_) and terminal (*U*_*T*_, *N*_*T*_) feature weights, the model can flexibly adjust how both uncertainty and novelty factor into the subjective utility of a given stimulus according to a linear trajectory across the task horizon.

As noted previously, participants expressed a growing distaste for uncertain options and a simultaneously increasing preference for novel options despite their inherent uncertainty, presenting a tension that demands further scrutiny. This pattern of response could emerge from a system that considers both novelty and uncertainty as pertinent stimulus features, but increases the drive to seek novelty at a rate that can outpace uncertainty aversion. However, we know of neither empirical evidence nor normative theory suggesting that novelty should be increasingly valued as a task approaches termination (and see Supplementary materials showing evidence that boredom or superstitions do not offer explanations consistent with the data). Alternatively, we reasoned that this pattern of behavior could emerge by way of an interaction between novelty and uncertainty processing in the brain; specifically, a system in which stimulus novelty interferes with uncertainty’s potency. In this framework, familiar options derive their subjective utility according to both reward history and an uncertainty bonus. Conversely, novel options are valued according to their optimistically initiated expected value alone, absent consideration of the uncertainty bonus. Under this scheme, as depicted in Figure 3A, an increasing reluctance to sample uncertain familiar options will push favor toward novel alternatives for which uncertainty is ignored, resulting in a growing propensity to sample novel options as the motivation for uncertainty-driven exploration diminishes.

**Figure 3:**
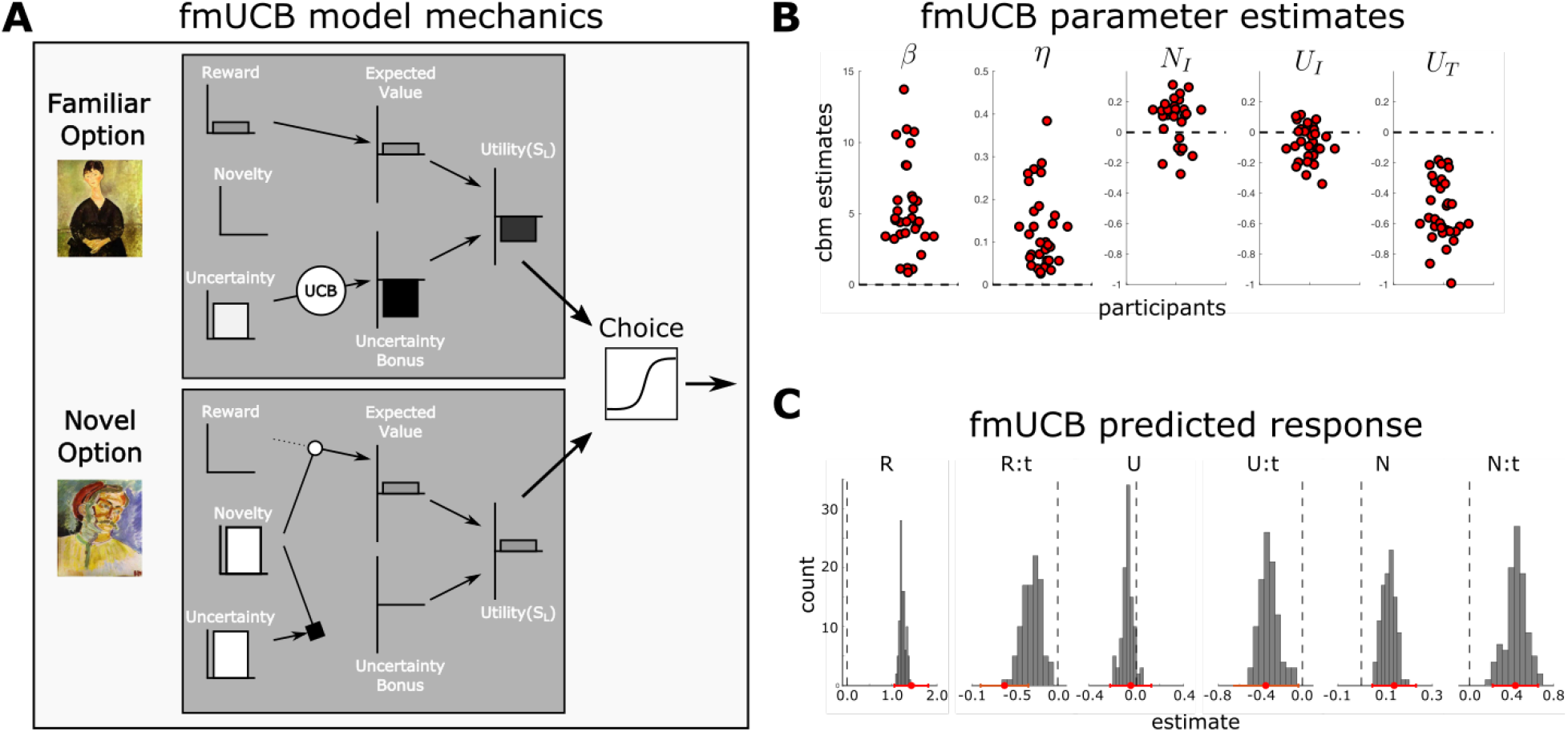
Computational modeling. **A)** An illustration of the mechanisms driving choice in the fmUCB model when novel and familiar options are offered near the end of a learning context. The familiar option’s negative utility combines the positive reward history coming from a previously rewarded encounter with a strong negative uncertainty bonus (due to the high uncertainty and proximity to the end of the task’s horizon). Despite not having been sampled, the novel option is endowed with a positive expected value while the otherwise aversive uncertainty is blocked, resulting in an positive stimulus utility. **B)** Parameter estimates for the fmUCB model (Softmax *β* governing choice determinacy, learning rate *η*, constant novelty shaping bias *N*_*I*_ = *N*_*T*_, uncertainty bonus intercept and terminal values *U*_*I*_, *U*_*T*_). **C)** Posterior predictive check depicting the degree to which the fmUCB model captures the response patterns highlighted by the computationally agnostic regression analysis. Histogram bars denote regression coefficient counts. behavioral regression coefficient and 95% confidence intervals are depicted as point and bars in red along the x-axis.

We formalized this hypothesis as a computational model that could explain the interaction between novelty and uncertainty to drive exploratory choice. We began with a baseline forgetful Bayesian reinforcement learning model that included two free parameters: a learning rate (*η*) that determined the rate of behavioral adaptation in response to observed outcomes; and a Softmax choice stochasticity parameter (*β*) which determined the degree to which choice relied on value. We augmented this baseline model to embody our hypothesized mechanisms of human exploration in what we label the familiarity modulated upper-confidence bound model (fmUCB). This model includes free parameters *U*_*I*_ and *U*_*T*_ to incorporate an uncertainty bonus that adapts to the task horizon, and *N*_*I*_ = *N*_*T*_ to facilitate a consistent value initiation bias throughout the task. Importantly, we address the tension between novelty and uncertainty by incorporating stimulus familiarity as a modulatory mechanism governing the uncertainty bonus, effectively blocking the influence of uncertainty when stimulus novelty is high.

We examined the degree to which the fmUCB model faithfully reproduced the response patterns observed in participants’ behavior by conducting a posterior predictive analysis. To do so we fit the model to behavior using the Computational Behavioral Modeling (cbm) toolkit [35]. We then exposed the model to the same sequence of trials observed by a participant from the fMRI task and had the model generate choices according to parameters optimized for that participant. This process was repeated for each participant to produce a synthetic data set that could be subjected to the same regression analysis applied to our human participant data (as illustrated in 2D). We repeated this process 100 times to generate a distribution of estimates, marginalizing over stochasticity in the model generated choices. Figure 3C illiterates the estimated effects identified in our participant group (in red along the x-axis), along side the distribution of effects generated by the model. The model can be seen to faithfully reproduce the patterns observed in the behavioral data, indicating that the fmUCB model does indeed capture patterns of interest in the behavioral data.

### 2.5 Model characteristics and comparisons on behavioral data

To test whether the fmUCB model offered a parsimonious explanation of participant’s behavioral data, we performed a model comparison in which we pitted it against alternative models. This alternative set included the baseline model described above (including *β* and *η* as free parameters), and a family of other models containing a full permutation of exploration-related variables absent the additional constraints of the familiarity gating mechanism, defined by a parameter set that includes initial and terminal novelty initiation bias (*N*_*I*_ and *N*_*T*_) as well as initial and terminal uncertainty bonus weights (*U*_*I*_, and *U*_*T*_). We performed model comparison on the behavioral data from the fMRI dataset which had sufficient numbers of trials per participant to enable model fitting and comparison. Using the cbm toolkit, this comparison showed that the fmUCB model offered the best explanation of the behavioral data given the set of models being considered with 96% exceedance probability.

As depicted in Figure 3B, the fmUCB model’s optimized parameters estimates exhibit a positive estimate of *N*_*I*_, indicating that option values are indeed optimistically initiated (t(31)=3.56, CI=[0.04, 0.14], p=0.001). Furthermore, participants also manifested a decreasing uncertainty bonus, consistent with the hypothesis that the uncertainty bonus encapsulates the prospective benefit of uncertainty reduction (*U*_*I*_ − *U*_*T*_ : t(31)=14.96, p<0.001, CI: [0.39, 0.52]).

We next considered critical details about how the fmUCB model could be implemented. Firstly, a reasonable alternative to optimistic initiation could involve novelty acting on the decision-making process as a separable linear bonus term that is integrated to form a subjective utility (as operationalized through the uncertainty bonus term). Thus, we sought to probe for evidence supportive of either an optimistic initiation or a linear bonus mechanism driving novelty seeking. Secondly, the fmUCB model does not specify the mechanism by which novelty dampens the uncertainty bonus. This pattern of response could emerge incidentally, where novel stimuli promote a decision before the bonus can be fully integrated into the subjective utility guiding choice, or alternatively, the neural processes engaged by novel stimuli could directly interfere with the processes required to compute the uncertainty bonus itself. Although these mechanisms are indistinguishable from a behavioral perspective (see below), the profile of the underlying computational variables as they evolve over time differs between the implementations. Given that these variables should be encoded in the brain according to the model variant that is actually being utilized at the neural level, we aimed to determine whether we could distinguish between these implementational forms using the fMRI data.

### 2.6 Neural correlates of subjective utility and preference

Before investigating the mechanistic implementation of the fmUCB model, we first sought to identify regions of the brain correlating with the model’s key variables. We focus first on identifying regions of the brain that are associated with the subjective utility of the stimuli being offered. To do so, we defined a GLM that included event onset regressors of zero duration (fixation, stimulus, response, and feedback), as well as a boxcar regressor lasting the trial’s duration. We augment this GLM to include the stimulus utility for both the selected and rejected options, which is defined as the summed expected value and uncertainty bonus as used by the fmUCB model to guide choice. We also include stimulus novelty and estimation uncertainty for both options as stimulus-locked parametric modulators to segregate signal associated with valuation, as well as the model’s reward prediction error as a feedback-locked parametric modulator.

We first probed the neural correlates associated with the selected option’s subjective utility (see Figure 4A, red). This analysis identified a positively correlated cluster comprising of vmPFC (peak voxel: x=-10, y=41, z=-10; t=6.4) and ventral striatum (peak voxel: x=5, y=16, z=-4; t=7.86), as well as a cluster spanning posterior cingulate cortex (peak voxel: x=-5, y=-51, z=10; t=5.44). We sought to further characterize these signals as contributing to valuation and/or the decision making process itself. Noting that the value of both options is correlated with their sum (selected + rejected), while a comparative decision making process is correlated with their difference (selected - rejected) [36], we constructed and analyzed the corresponding contrasts. The summed subjective utility contrast tracking value ascribed to either option identified a cluster extending from vmPFC (peak voxel: x=-8, y=36, z=-14; t=5.3) to ventral striatum (peak voxel: x=12, y=6, z=-7; t=5.39), and a small cluster in posterior cingulate cortex (peak voxel: x=-5, y=-41, z=40; t=4.36) (Figure 4B red). Focusing next on the decision making process, we identified a cluster in vmPFC (peak voxel: x=-10, y=46, z=-12; t=4.91) that was positively correlated with the relative difference in subjective utility (selected-rejected; Figure 4C red). Consistent with numerous previous studies [28, 27], these results highlight the vmPFC as playing a central role in the valuation and decision making process.

**Figure 4:**
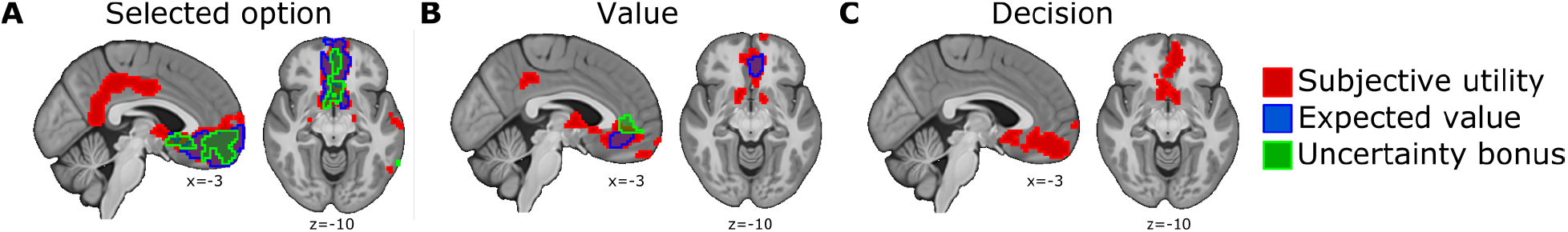
Neural correlates of computational variables guiding choice **A)** Neural correlates associated with the chosen option. Clusters including posterior cingulate gyrus, and a large cluster extending from vmPFC to ventral striatum were positively correlated with the subjective utility of the chosen stimulus (red). A cluster including vmPFC and ventral striatum was positively correlated with the expected value (blue), while a cluster incorporating vmPFC was positively correlated with the uncertainty bonus (green). **B)** Value contrasts (selected + rejected option value) exposed a cluster extending from vmPFC to ventral striatum as positively correlated with the mean subjective utility of both options (red). A cluster in vmPFC was positively correlated with expected value (blue), while a cluster in mPFC was correlated with the uncertainty bonus (green). **C)** Decision contrasts (selected - rejected) identified a significant cluster associated with subjective utility extending from vmPFC to ventral striatum. Whole-brain maps for signal of interest were tested with a cluster-forming threshold of *P* < 0.001 uncorrected, followed by cluster-level FWE correction at P<0.05.

The fmUCB model derives the subjective utility for each option as the sum of an optimistically initiated expected value and a familiarity modulated uncertainty bonus. We sought to determine whether these signals were represented in the brain, and if so, how they come to influence the decision making process. In pursuit of this, we specified a second GLM in which the subjective utility was decomposed into its constituent expected value and uncertainty bonus terms for both the selected and rejected option. An analysis of BOLD signal change positively correlated with the selected option stressed a prominent role for vmPFC in tracking stimulus features pertinent to value and decision making (see Figure 4A blue and green), with clusters positively correlated with novelty-biased expected value (peak voxel: x=-8, y=36, z=-10; t=6.62) as well as a largely overlapping cluster positively correlated with the uncertainty bonus (peak voxel: x=-2, y=51, z=-4; t=5.1). We then repeated the valuation (selected+rejected) and decision (selected-rejected) contrast analyses for both expected value and the uncertainty bonus. The valuation contrast exposed a cluster in vmPFC that positively correlated with optimistically initiated expected values (peak voxel: x=-8, y=36, z=-14; t=4.89, Figure 4B blue), while a slightly more dorsal cluster in medial PFC (peak voxel: x=8, y=42, z=8; t=4.53, Figure 4B green), was positively correlated with the uncertainty bonus. No regions were found to correlate with the relative difference between the option’s expected values or between their uncertainty bonus terms. These results suggest that stimulus features themselves are not directly compared; rather, an integrated subjective utility is formed and used to guide choice.

### 2.7 fMRI evidence for optimistic initiation

As noted previously, there are two different routes by which novelty could be incorporated to guide choice. Besides the optimistic initiation mechanism described earlier, an alternative would involve the brain employing a separable novelty bonus term applied at the time of decision, which we call a novelty bonus mechanism. A simulation-based analysis of model confusability showed that these two candidate mechanisms are not identifiable using behavior alone as the generative mechanism can only be correctly identified 54% of the time (see Methods for details). Crucially, although the choices motivated by both mechanisms are roughly equivalent, they differ with respect to the expected value ascribed to novel options and their subsequent RPEs when chosen. Thus, while not behaviorally distinguishable, we aimed to determine whether fMRI activity could be utilized to differentiate between these two implementations.

Exploiting the fact that optimistically initiated values result in distorted reward prediction errors when outcomes are observed, we first examined whether signal change in ventral striatum reflects the bias predicted by optimistic initiation as previously reported by [6]. We extracted two reward prediction error signals from the fmUCB model’s time-course; the first RPE (*δ*) represents the signal generated by the fmUCB model that includes the optimistic initiation component, and the second is the RPE as would be computed absent any effect of novelty on value expectations (*δ*_−*N*_). From this we defined a regressor representing the novelty component of the RPE as *δ*_*N*_ = *δ* −*δ*_−*N*_. Consistent with previous findings [20, 37], we found a strong correlation between the standard RPE (*δ*_−*N*_) and activation in ventral striatum (see Figure 5A). Importantly, voxels in right ventral striatum also correlated with the novelty biased component of the RPE encoded by *δ*_*N*_ above and beyond the correlation found with the basic RPE signal (peak voxel: x=10, y=16, z=-9; t=3.02, p<0.005 height threshold; p=0.05 SVC). These findings support the presence of a novelty biased RPE signal, consistent with (and overlapping spatially with) previous findings reported by [6].

**Figure 5:**
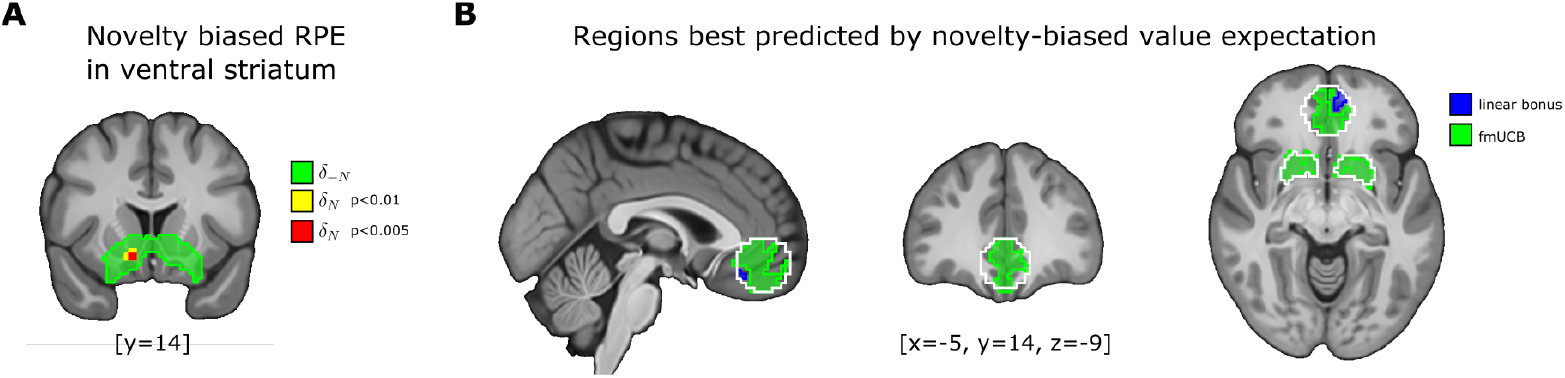
Representation of optimistic initiation in vmPFC and ventral striatum **A)** Standard reward prediction errors are associated with robust activity in ventral striatum (green). The additional component of the RPE attributed to optimistic initiation was also identified in ventral striatum. **B)** Signal change in vmPFC and ventral striatum is better explained by optimistic initiation than a linear bonus mechanism. Voxel-wise model comparison plots report voxels exceedance probability > 0.9 favoring the fmUCB model (green) or the linear bonus model (blue). Mask encompassing vmPFC and ventral striatum bordered in white.

Pursuing this line of investigation further, we compared the variance explained by the optimistic initiation and linear novelty bonus mechanisms in regions of interest using Bayesian model comparison. We identified the vmPFC as a region of interest associated with valuation and choice [27, 28], defining a 15mm radius sphere centred on peak voxel coordinates identified by [27]. We also probed ventral striatum as a second ROI given its association with reward prediction errors as found above [6, 37]. As depicted in Figure 5B, a comparison of the variance explained by the expected value associated with the chosen option and the subsequent RPE as predicted by both mechanisms shows that 421 voxels in vmPFC and 357 voxels in ventral striatum were best explained by the optimistic initiation mechanism (exceedance threshold >= 0.9, cluster size >=10). Conversely, only 94 voxels were best explained by the linear bonus mechanism. Therefore, our findings suggest that the fmUCB model operationalized in terms of optimistically initiated reward value expectations provides a better overall account for expected value and RPE signals in both vmPFC and ventral striatum than does the linear bonus mechanism. It should be noted that value and prediction error signals derived from both the optimistic initiation and linear bonus implementations of the fmUCB model yield significant effects in overlapping brain areas at the whole brain level, and these are already encompassed by our ROIs. Thus, our model comparison conclusions are not merely an artifact of the ROIs selected because these ROIs capture the key regions in which statistically significant fMRI correlates of these variables are present according to either implementation (see Supplementary Figure 1 for details).

### 2.8 Novelty disrupts the computation of the uncertainty bonus

We also wanted to determine precisely how uncertainty and novelty interact within the human brain. The fmUCB model flexibly adapts the uncertainty bonus according to the task’s horizon, growing increasingly unwilling to probe uncertain familiar options toward the end of a learning context (see Figure 3D-E). The model also expresses a growing preference for novel options despite their inherent uncertainty, which is accommodated through a familiarity modulated uncertainty bonus. Here, we query the fMRI data for patterns consistent with two plausible candidate installations of the model’s uncertainty bonus.

As previously outlined, this pattern of response could emerge incidentally as a result of novelty promoting a choice before the uncertainty bonus could be fully integrated into the subjective utility. Alternatively, the neural processes required to compute the uncertainty bonus may be directly obstructed or commandeered by processes engaged with novel stimuli. These two competing implementations cannot not be distinguished on the basis of the behavioral data alone as they yield equivalent choice behavior. Therefore, we sought to utilize the constraints imposed by their neural implementation to differentiate between them. Specifically, direct interference predicts the absence of uncertainty bonus signals associated with novel stimuli, while incidental choice induction implies that the cognitive processes responsible for computing the uncertainty bonus ought to be present despite their inability to guide behavior.

We again used a model comparison approach to determine which of these implementations better describe the pattern of activity in the brain. Our previous analyses show that the uncertainty bonus is reflected in mPFC (see Figure 4B), highlighting this as a target ROI within which to evaluate the predictions of the two mechanisms. We first defined a GLM that included event onset regressors of zero duration (fixation, stimulus, response, and feedback), as well as a boxcar regressor lasting the trial’s duration. Reflecting the premise that uncertainty bonus signals should emerge for all stimuli regardless of novelty, the incidental mechanism was modelled in a GLM that included expected value, novelty and uncertainty for both options offered, as well as their trial-weighted uncertainty bonus terms absent familiarity modulation as parametric modulators locked to stimulus onset. We compared this GLM to one in which the uncertainty bonus signals were dampened by novelty as used by the fmUCB model and reported in Figure 4B. As illustrated in Figure 6A, a cluster of 80 voxels was found to be better represented by the familiarity modulated bonus term, whereas no voxels favored the uncertainty bonus absent familiarity modulation (exceedance probability > 0.9, cluster threshold = 10). Importantly, although no significant clusters were found to correlate with the unmodulated uncertainty bonus, peak voxels were identified in this ROI at an extremely liberal threshold, demonstrating the suitabiliy of the ROI for comparison (see Supplementary Figure 2). This suggests that the brain’s uncertainty bonus is diminished when processing novel stimuli, and is consistent with a mechanism in which novelty actively inhibits the processes through which uncertainty directed decision making is guided.

**Figure 6:**
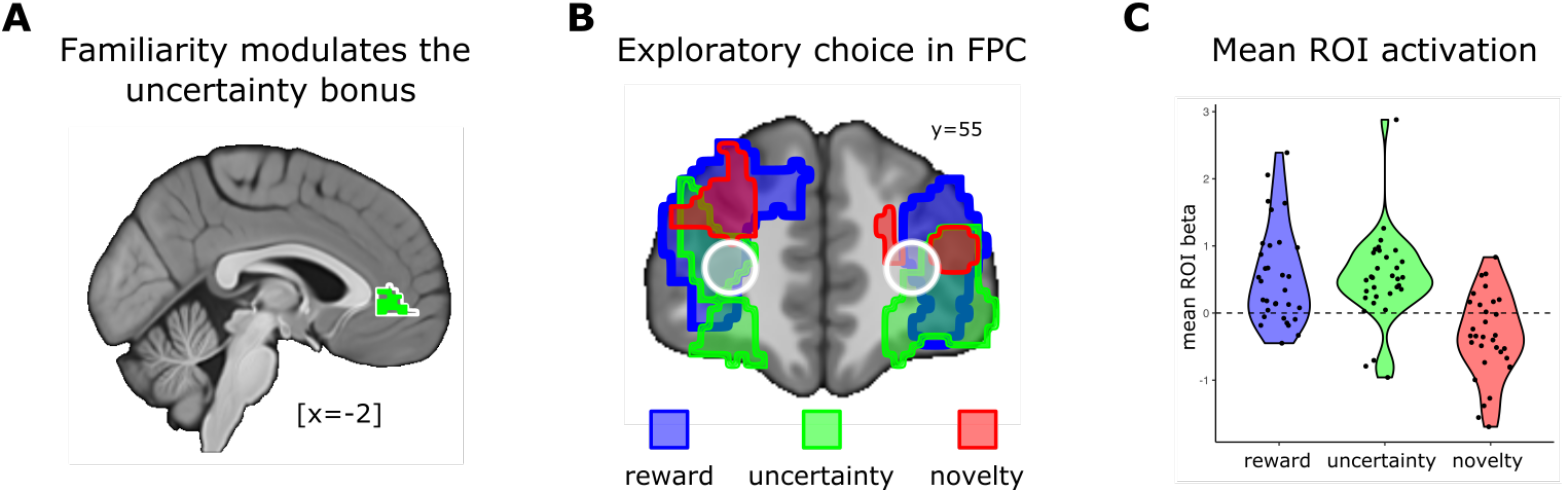
Effects of novelty on the uncertainty bonus and choice. **A)** Model comparison in mPFC supports a novelty-induced interference implementation of the familiarity modulated uncertainty bonus. Voxel-wise model comparison report the exceedance probability favoring the explicit interference implementation of the fmUCB model. Masked brain plots report voxels favoring the explicit fmUCB model exceeding threshold (exceedance probability > 0.9) in green, (mask encompassing mPFC bordered in white). **B)** Contrasts showing significant clusters in bi-lateral FPC positively correlated with choosing the lower valued option (blue), the more uncertain option (green), and negatively correlated with choosing the more novel option (red). Brain maps reported exceed a cluster-forming threshold of *P* < 0.001 uncorrected, followed by cluster-level FWE correction at P<0.05. **C)** ROI analysis of mean beta estimates in ROIs situated at bilateral FPC (white bordered circles) taken from [20] according to choice.

Inquiring further into this idea that novelty hampers the brain’s application of the uncertainty bonus, we probed for regions expressing divergent neural correlates tied to novelty and uncertainty processing. Ultimately, the computational variables and cognitive processes under investigation produce a behavioral choice. Here we leverage the fact that our experimental design offered options that varied in terms of expected value, estimation uncertainty and novelty to probe the activation patterns across these dimensions when choice strayed from reward exploitation.

We constructed a GLM that included three response-locked parametric regressors indicating whether or not the option with lower expected value was selected (random exploration), if the option with higher estimation uncertainty was sampled (uncertainty directed exploration), and whether or not a novel option was chosen (novelty driven exploration). As illustrated in Figure 6B, this analysis revealed differential bi-lateral engagement in FPC for each class of choice. We identified bi-lateral clusters in FPC that showed elevated activation when either the lower-valued option was chosen (rFPC: x=32,y=51,z=20 t=5.94; lFPC: x=-27,y=51,z=15 t=5.4), or when the option with higher uncertainty was sampled (rFPC: x=25,y=56,z=-12 t=5.96; lFPC: x=-25,y=64,z=0 t=4.49). In contrast, we observed a significant bi-lateral reduction in FPC activity when novel stimuli were chosen (rFPC: x=22,y=51,z=23 t=4.91; lFPC: x=-20,y=46.5,z=13 t=5.02). These effects were corroborated by an ROI analysis centered on FPC coordinates previously reported to be associated with exploration [20] (Figure 6B-C), with both value (t(31)=4.3, p<0.01) and uncertainty (t(31)=4.3, p<0.01) driven exploration associated with greater FPC activation, while sampling novel options elicited a significantly reduced activation in those same regions (t(31)=-3.2, p<0.01).

These patterns of activation associated with non-exploitative choice show that while decisions to explore familiar options preferentially engage FPC, a region that has been repeatedly associated with exploration [20, 24, 8, 38], choice targeting novel alternatives does not. Further to this point, sampling novel options corresponds with depressed activation in these same regions, suggesting that the exploratory contributions fostered by FPC are impeded by stimulus novelty.

## 3 Discussion

In this study we investigate how the human brain balances a fundamental tension between exploration and exploitation. Previous studies have characterized a range of normative and heuristic solutions to the problem [20, 7, 8, 21, 10, 6, 9, 19, 29]; however, none have simultaneously probed the interactions between novelty and uncertainty, two key variables we have shown to co-exist, interact and influence choice behavior during exploration/exploitation decisions. Here, we show that humans adaptively evaluate the potential benefit of reducing uncertainty, preferentially targeting uncertain options when new information can be exploited in the future. In contrast, and despite their inherent uncertainty, novel options were pursued irrespective of the task horizon, and grew increasingly attractive as task termination approached. Notably, these findings were also replicated in a larger behavioral sample, demonstrating generalizability across populations.

Using a computational model of choice constrained by neural data, we demonstrate how this apparent tension between uncertainty-guided choice and novelty seeking can be resolved. We found that choice was biased by an adaptive uncertainty bonus that reflects the prospective value of information. At the start of each learning context, participants were more likely to be drawn toward uncertain options; however, as the opportunity to leverage newly gained information diminished so too did the willingness to explore uncertain alternatives. This is a reasonable course of action from a theoretical point of view, where the value of information is rooted in the rewarding advantages it grants in the future [2], and mirrors previously reported effects of task horizon linking uncertainty driven exploration to the prospective value of information [7].

Consistent with the hypothesis that novelty seeking reflects a mechanism encouraging exploration though the exploitation of corrupted reward expectations, participants did not sway from seeking novel options throughout the task. And indeed, our results show that signal change in ventral striatum, a region that has previously been associated with novelty biased reward prediction errors [6], and vmPFC, a region associated with both value guided choice [27] and outcome monitoring [28], were both best accounted for by a model that included an optimistic initiation mechanism (see Figure 5B). Furthermore, signal change in ventral striatum associated with reward prediction errors was shown to include an additional novelty bias component, replicating findings reported previously [6] (Figure 5A).

Contrary to normative theory, where the value of exploration stems from the prospects of greater reward in the future [2], we also found that participants exhibited an increasing tendency to sample novel options as the task horizon approached its end, a pattern that optimistic initiation alone could not explain. Our analyses show that, in addition to inflating an option’s expected value, novelty also diminished the influence of uncertainty. Model comparison demonstrated that the fmUCB model, which included a mechanism through which stimulus familiarity modulated the influence of the uncertainty bonus, fit the behavioral data best. Furthermore, we found that signal change in mPFC, where correlates of the uncertainty bonus and subjective utility were localized, was better explained by a familiarity-modulated uncertainty bonus than by a model that applied the uncertainty bonus regardless of stimulus novelty (see Figure 6A). This, we argue, is consistent with an antagonistic interaction between novelty and uncertainty processing as opposed to an indirect relationship in which novelty promotes response initiation before the uncertainty bonus can be integrated into the decision making process. Thus, we propose that the increasing appetite for novel options reflects two separable processes that act to promote exploration: firstly, inflated reward expectations guide the brain’s exploitative circuitry towards sampling novel options, and second; inhibited uncertainty processing diminishes the otherwise aversive nature of the unknown when new information has low prospective utility.

The present study also has novel methodological implications that go beyond the specific research question. It provides a clear demonstration of how neural evidence can be used to constrain and arbitrate between different computational mechanisms in a way that sometimes cannot be achieved using behavioral measures alone. This demonstration is pertinent to a persistent debate in the psychological literature about the utility (or absence thereof) of neuroscience measures above and beyond behavioral measures for advancing theoretical understanding of cognitive function. While the behavioral patterns we observed on the task allowed us to restrict the space of possible computational models, behavior could not distinguish between several competing implementations of the model describing how stimulus novelty influenced choice. However, variants of the model’s instantiation relied on distinct internal signals, allowing us to compare patterns of brain activity that would be expected for different model components including the representation of expected value, prediction errors and uncertainty bonus signals. Utilizing a model comparison approach on fMRI data, it was possible for us to obtain evidence in support of one particular model structure, thereby validating the importance of using brain measures alongside behavioral ones to advance theoretical understanding of cognitive processes [39, 40, 41, 17].

Our results also build on a wealth of previous work exposing the role of the vmPFC in learning and decision making. In contrast to tasks in which decisions are either purely exploitative or exploratory [10, 11], or where exploration was prompted by latent change-points indicating the need for a new strategy [29, 24], our task was explicitly designed to encourage choice that simultaneously considered both reward and information gain as prescribed by normative theory [2]. Previous investigations of choice under uncertainty have shown that vmPFC tracks both the probability of choice [20], and the relative difference between the selected and rejected values [24], implicating vmPFC in both valuation and decision making [27]. In line with these reports, we found that both the subjective utility of the options under consideration and their relative difference was robustly represented by vmPFC (Figure 4B-C).

Recent studies have suggested that vmPFC contributes to the regulation of exploration and exploration by monitoring outcomes, signalling the degree to which predictions are being met (or not) [29], while others have reported vmPFC signal consistent with the value ascribed to information seeking, be it a positive value when information can be leveraged in the future, or a negative value when it cannot [11]. Our results build on these findings, showing that vmPFC plays a role in regulating the trade-off between exploration and exploitation by integrating feature values pertinent to each. We identified signal consistent with the expected value of the options offered in vmPFC, and an uncertainty bonus guiding exploration in mFPC. Notably, while we observed a robust signal in vmPFC reflecting the relative difference in subjective utility, we did not observe signal consistent with the difference in feature values. This suggests that the expected value of reward and the expected value of information gained by exploration are first integrated to form subjective stimulus value from which choice may be directed. This is consistent with previous work arguing that vmPFC acts as a value integration hub where disparate stimulus features are weighted according to current goals, situating vmPFC as integral to goal-directed action selection [42, 43, 44].

Previous research examining the neural correlates of human exploration have implicated FPC; however, the nature of its role remains elusive. One of the first studies using fMRI to probe exploration in humans found that this region was engaged when participants opted to forgo the most rewarding option to sample a lower valued alternative [20], implicating FPC in random exploration. Subsequent work has associated FPC in tracking the relative uncertainty of the available options [8, 23], while others have proposed a functional link between FPC and IPS as a means of tracking and switching to an alternative course of action [24]. Consistent with these findings, we found that FPC was preferentially engaged when participants decided to sample either the lower-valued or more uncertain option (Figure 6B-C). Thus, our data highlight the role of FPC in both directed and random exploration, suggesting that this region may be critical for motivating a diverse range exploratory strategies.

In contrast, we also found that FPC was suppressed when novel options were chosen. This, we argue, offers further support for the hypothesis that the brain does not frame novelty seeking as exploration per se; but as value exploitation. Furthermore, noting the uncertainty inherent to novel options, these findings may reflect the active suppression of signals pertinent to uncertainty guided choice. This intriguing possibility could potentially be tested in follow up work leveraging connectivity analyses and neural stimulation over the FPC.

Questions remain as to what benefit this particular configuration offers, with stimulus novelty parasitizing the brain’s reward circuitry to promote exploration while uncertainty-directed sampling relies on separable integrated feature values. We speculate that this offers a parsimonious decomposition of an otherwise intractable decision problem. Inferring the prospective benefit of strategically reducing uncertainty can offer significant benefit, particularly in volatile environments where reward contingencies and goals can change rapidly. However, the computational demands of this approach are high, and in unfamiliar circumstances, most likely nothing more than guess work. We suggest that by diverting computational resources away from ill suited strategies in favor of more computationally efficient heuristics like optimistic initiation, the brain can begin to bridge the gap from a computational intractable scenario toward a manageable landscape where more adaptive strategies like uncertainty-driven exploration can be beneficially applied.

To conclude, in this study we offer further insight into human strategies for balancing the explore/exploit trade-off and their neural roots. By systematically decoupling stimulus novelty and uncertainty, and by leveraging neural data to constrain models of human learning and decision making, we show that human exploration simultaneously targets different stimulus features using distinct strategies with potentially conflicting preferences. How the brain resolves these tensions has a significant impact on behavior, highlighting potential avenues through which dysregulation of the balance between exploring new alternatives vs staying the course may be investigated.

## 4 Materials and Methods

### 4.1 Participants

#### 4.1.1 fMRI study

We recruited 33 participants from the Pasadena community to take park in our study. One participant was removed from the sample due to excess movement and poor performance (sleeping in the scanner), leaving a sample of 32 human participants (13 female). All participants were English speakers, had normal/corrected-to-normal vision, and had no history of neurological or psychiatric disease. Participants were paid a $40 base-rate plus a performance bonus ranging from $5-$15. The study was approved by the Caltech IRB and participants gave their informed consent to take part in the study.

#### 4.1.2 Behavioral replication study

The replication study included 79 participants (48 female) from the Toronto community. All participants were required to be fluent English speakers and have normal or corrected-to-normal vision. 77 participants reported no prior history of psychiatric or neurological disease, but 2 additional participants reported a prior history of psychiatric illness. Those two participants are still included in the reported analyses because omitting them made no substantive difference to the results. Participants were paid a base rate of $10 per hour plus a performance bonus up to $20. All participants gave their informed consent to participate in the study in accordance with the Research Ethics Board at the University of Toronto.

### 4.2 Experimental design

#### 4.2.1 fMRI study

Participants performed 20 blocks of a novel finite horizon multi-armed bandit task designed to expose the impact of reward, uncertainty and novelty in balancing the explore/exploit trade-off while undergoing four consecutive 15-minute fMRI sessions (5 blocks per session). On each trial, participants were asked to choose between two slot machines. Having selected one of the machines, they were informed of the machine’s payout, either $1US (win) or $0US (loss). Participants were instructed that each block would consist 18 to 23 trials, and to encourage balanced attention and motivation throughout the task, they were informed that one block would be selected at random at the end of the experiment and they would be awarded the earning collected during that block as a cash bonus.

Each block was structured to include five visually unique and identifiable slot machines, three of which used familiar stimuli that had been seen during previous blocks, while the remaining two used novel stimuli that had not yet been shown (43 stimuli in total). The five slot machines that could be offered in a given block were each associated with a fixed probability winning, which was sampled from a linearly spaced range of [0.2 - 0.8]. Participants were informed that each slot machine had a fixed probability of winning within a block, but all machines were re-set at the start of each block, and as such, anything learned in previous blocks would not apply in blocks to come. Trials were structured such that the two slot machines offered on each trial varied in terms of the number of previous exposures (novelty manipulation), as well as the number of times they’d been sampled in the current block (uncertainty manipulation), allowing us to to systematically examine the influence of novelty and uncertainty across the horizon of trials within a given block.

Following the slot machine task, participants performed a recognition test designed to probe their recall of which machines they had observed. They were asked to label 86 stimuli as ‘old’ or ‘new’, half of which had been used during the multi-armed bandit task, and half of which were not. Participants exhibited exceptional performance on the memory probe task, with mean accuracy of 89% (min 77%, max 97%), indicating the efficacy of the novelty manipulation.

#### 4.2.2 Behavioral replication study

The replication study consisted of an adapted version of the task described in the preceding section, with modifications described here. Prior to the start of the task, participants were instructed that they would be playing for points that would then be converted to a real monetary bonus up to a maximum of $20cnd. Because this version of the task included situations of monetary loss in affective conditions instantiated after an initial baseline condition, participants were initially endowed with a starting sum of 1200 pts, with machines paying out 50 pts for a win and 0 pts otherwise. The baseline condition mirrors the design of the fMRI study; the affective manipulations only start after the baseline condition is complete and will be reported elsewhere.

Participants completed 6 blocks of the baseline condition consisting of 23 trials each. On each trial, participants were asked to chose between two slot machine from a set of six, each of which was associated with a fixed probability of winning, either sampled form a linearly spaced range of [0.2, 0.8] or from the set comprising [0.2, 0.44, 0.48, 0.52, 0.56, 0.6]. The assigned set of fixed win probabilities for a given block was chosen randomly, and participants were similarly instructed that the machines re-set at the start of each block. The structure of the novelty and uncertainty manipulations followed that reported in the fMRI study, though with two novel and four familiar stimuli. Two familiar stimuli were presented for the first 2 trials, with the first novel and third familiar stimuli introduced between trials 3-5. The remainder of the set (second novel and fourth familiar stimuli) were introduced between trial 8-19 in a pseudo-randomised manner. Relevant for the affective manipulation but independent of the primary multi-armed bandit task reported here, participants were probed with a subjective mood rating scale in 2 of the 6 blocks; these ratings are not analyzed further here.

As with the fMRI study, participants performed a recognition memory task following the multi-armed bandit task. In the replication sample participants similarly exhibited good recognition memory accuracy (mean = 82%, s.d. = 12%)

### 4.3 fMRI data acquisition

Imaging data was collected at the Caltech Brain Imaging Center (Pasadena, CA) using a 3T Siemens Magneto TrioTim scanner using a 32-channel radio frequency coil. Functional scans were acquired using multiband acceleration of 4, 56 slices, voxel size = 2.5mm isotropic, TR = 1,000 ms, TE = 30 ms, FA = 60°, FOV = 200mm × 200mm. T1 and T2 weighted anatomical high-resolution scans were collected with 0.9mm isotropic resolution following the functional scans collected during task play.

### 4.4 fMRI data preprocessing and analysis

Data was preprocessed using a standard pipeline for preprocessing of multiband data. Using FSL [45], images were brain extracted, and denoised using ICA component removal, where components were extracted using FSL’s Melodic, and classified into signal or noise with a classifier trained on independent datasets. Functional data was then aligned, high-pass filtered (100 s threshold), and unwarped. T2 images were aligned to T1 images with FSL FLIRT, then both were normalized to standard space using ANTs (using CIT168 high resolution T1 and T2 templates [46, 47]). Functional data was co-registered to anatomical images using FSL’s FLIRT, then registered to the normalized T2 using ANTs. Finally, the functional data was smoothed using a 8mm FWHM Gaussian kernel. GLMs were specified using default specifications in SPM 12 [48]. The details of each first level GLM are provided in the main text. Second level T-maps were constructed by combining each subject’s first level contrasts with the standard summary statistics approach to random-effects analysis implemented in SPM. Statistic images were assessed for cluster-wise significance using a cluster-defining voxel threshold of p<0.001 and brain-volume cluster corrected threshold of FWE< 0.05.

### 4.5 Regions of interest and small volume correction

The 15mm radius sphere in vmPFC used for analyses reported in Figure 5B centered on peak vmPFC voxel reported in a meta analysis of the neural correlates of decision making and subjective utility [x=-2, y=40, z=-6] [27]. Center coordinates for left ([x=-14, y=10, z=-6]) and right ([x=14, y=10, z=-6]) ventral striatum were defined using coordinates from NeuroSynth, while the ROI was defined as the union of 15mm spheres centered at both left and right coordinates in union with voxels labeled as comprising putamen or accumbens according to the Harvard-Oxford sub-cortical structural atlas. FPC ROIs were defined using 5mm radius spheres centered on peak voxels in left and right FPC as reported by [20]. Spherical small volume corrected analysis applied to novelty-biased reward prediction errors reported in Figure 5A used a 9mm sphere centered on coordinates [x=18, y=16, z=-10] as previously reported by [6].

### 4.6 Statistical analyses

Behavioral data was analyzed using mixed-effects logistic regression for a descriptive characterization of task performance (using lme4 package in R). We define each option’s expected value (*E*[*S*_*i*_]) as the mean of a Beta distribution specified according to the number of wins and losses observed within the current block of trials (Beta[*α* =number of wins + 1; *β* =number of losses + 1]). Uncertainty (*U*[*S*_*i*_]) was defined as the variance of the same Beta distribution. We define stimulus novelty (*N* [*S*_*i*_]) as the variance of a Beta distribution specified according to the number of times a particular stimulus had been observed across the entire experiment (Beta[*α* =number of exposures + 1, *β*=1]). Probability of selecting the option presented on the left (*a*_*t*_ = *L*)for each trial *t* was modelled as :

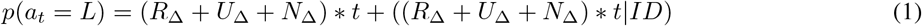

where *R*_∆_ = *E*[*S*_*L*_] − *E*[*S*_*R*_] denotes the reward differential in favor of the left option, while *U*_∆_ = *U* [*S*_*L*_] − *U* [*S*_*R*_], and *N*_∆_ = *N* [*S*_*L*_] − *N* [*S*_*R*_] reflect the difference in uncertainty and novelty respectively. We include trial number *t* as an interaction term to model changes in feature influence across the block horizon, with random intercept and slopes estimated all terms for each participant ID.

### 4.7 Computational modeling

We characterize the computational mechanisms balancing the trade-off between exploration and exploitation using a forgetful Bayesian model of choice, augmented with an uncertainty bonus and optimistic value initiation. The subjective utility derived for each stimulus *s*_*i*_ is defined as:

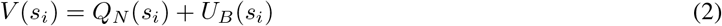

where *U*_*B*_(*s*_*i*_) is the uncertainty bonus added to the optimistically initiated expected value, *Q*_*N*_(*s*_*i*_). The probability of selecting either the option presented on the left (*s*_*l*_) or right (*s*_*r*_) was derived using a Softmax function, meaning choice was a function of both random and uncertainty-directed exploration:

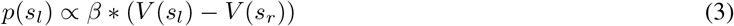

where *β* is the Softmax parameter controlling the degree to which choice was determined by value.

The forgetful Bayesian reinforcement learning agent maintains a representation of each slot machine in a given block of trials as a Beta distribution. This distribution was defined according to a recency weighted integration of observed outcomes:

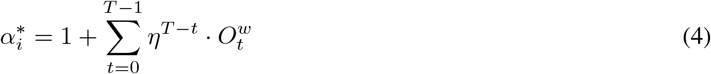

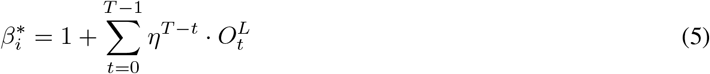

where *T* denotes the current trial within the block, and 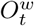 and 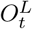 are binary flags noting whether or not the observed outcome on trial *t* was a win or loss respectively. Thus, *η* operates as a learning rate, controlling the rate at which past outcomes are down-weighted in favor of more recent outcomes. Each option’s expected value was derived as the mean of this distribution:

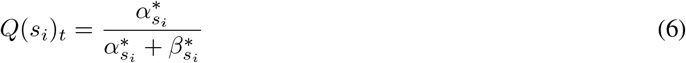

while uncertainty was derived as the variance of the same distribution:

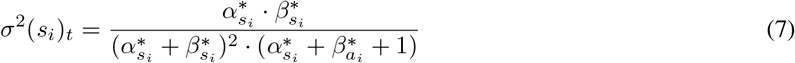

Optimistic initiation was integrated into the model by way of inflating the hyper-parameters describing the expected value 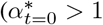 for a novelty seeking bias, and 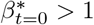 for a novelty avoidance bias).

Both the novelty-induced optimistic initiation bias and the uncertainty bonus were subject to dynamic weighting terms defined according to the block’s horizon. On each trial, weighting terms 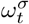 and 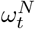 are applied to the uncertainty bonus and optimistic initiation values respectively, where weights are defined as a linear function of the current block’s trial number:

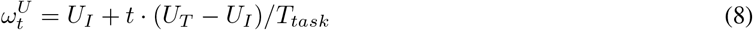

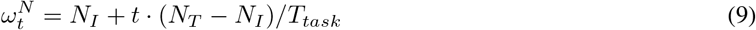

where *T*_*task*_ denotes the task horizon, *U*_*I*_ and *N*_*I*_ denote intercepts for uncertainty and novelty at the start of each block, while *U*_*T*_ and *N*_*T*_ denote weights at the end of each block. Thus, 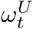 and 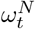 reflect linear trajectories across the task horizon.

We define the fmUCB model as a resolution of the tension between novelty seeking and uncertainty aversion. We embody this mechanisms by way of a familiarity modulated uncertainty bonus. Stimulus familiarity was defined according to the normalized variance of a Beta distribution defined according to hyper parameters (Beta[*α*(*s*_*i*_) =number of observations + 1, and *β* = 1]), or specifically:

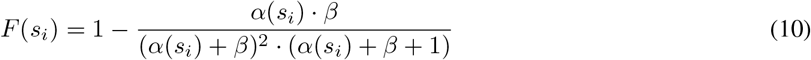

in which the variance terms is scaled to range between [0,1]. We then augment the uncertainty bonus to also reflect stimulus familiarity:

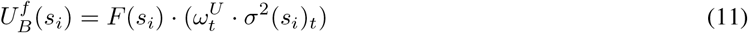

Lastly, in contrast to novelty biasing the expected value, we also test a mechanism in which novelty is factored into the decision making process as its own bonus feature, referred to as the linear bonus model. In this model the optimistic initiation mechanism was removed from the fmUCB model, and augment the subjective utility to include a novelty bonus term:

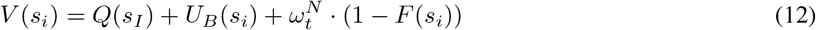

### 4.8 Parameter estimation and model comparison

All model parameters were estimated using the Computational Behavioral Modeling (cbm) toolkit [35]. The toolkit relies on normally distributed parameters, meaning some model parameters need to be transformed to be sensible. Novelty and uncertainty weighting terms (*N*_*I*_, *N*_*T*_, *U*_*I*_, *U*_*T*_) all remained normally distributed (no transformation). The Softmax inverse temperate was constrained to range between [0 ≤ *β* ≤ 20], while the learning rate was constrained to range between [0*≤ η ≤* 1] using a sigmoid function. Initially, we used an exponential function to transform the Softmax parameter (*β* = *exp*(*p*_1_)); however, this resulted in convergence/halting problems as small changes in *p*_1_ could lead to very large or very small changes in *β* depending on the sign of *p*_1_. Noting that all estimates were within a [0:20] range when using the exponential transformation, we opted to use a sigmoid function as it offered a smoother and more balanced transformation.

Model comparison was also relied on the cbm toolkit [35]. First-level estimates were computed for each individual and model using common priors (*𝒩*(*µ* = 0, *σ*^2^ = 6.25)), which were then used to inform second-level fits and simultaneous model comparison. This process treats model comparison as a random effect (i.e. different models might better represent different participant), while also taking advantage of hierarchical parameter estimation which relies on empirical as opposed to prescribed priors.

## 5 Supplementary Materials

### 5.1 Learning contexts were treated independently

Our experimental design decoupled estimation uncertainty and stimulus novelty by framing each block of trials as a unique learning context, and as such, the value associated with a stimulus sampled during previous contexts should not be applied to the current context in play. Participants were explicitly informed of this, and told that although they may see the same slot machines in different casinos, each casino has programmed the winning probabilities for each machine differently. Here we demonstrate that participant behavior faithfully reflects de novo learning in each context, and thus our assumption that the expected value and uncertainty associated with stimuli encountered during previous contexts can be appropriately modeled using an unbiased prior at the start of a learning block (Beta[*α* = *β* = 1]).

We first probed for behavioral signatures of value carry over from previous contexts by augmenting the fmUCB model to accommodate value carry-over from previously learned stimulus values. To do so, we define the Beta distribution describing stimulus *i*’s expected value according to:

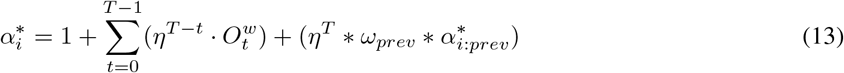

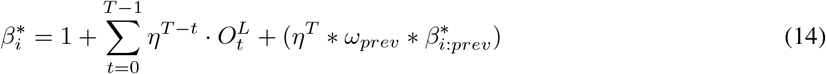

Here, the expected value is derived according to wins and losses observed in the current context up to the current trial *T*, where *η* is a free parameter regulating the rate at which previous outcomes observed in the current context are down-weighted in favor of more recent observations (just as the ‘forgetting rate’ defined in the fmUCB model). The Beta distribution’s parameters for stimulus *i* also include some proportion of the value associated with that stimulus in the most recent previous context (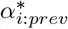 and 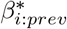), where *ω*_*prev*_ governs the proportion of value carry over, and *η*^*T*^ down-weights the initialization value as trials proceed in order to accommodate temporal effects of learning.

We fit and submitted the fmUCB model, a model with *ω*_*prev*_ = 1, and a model with *ω*_*prev*_ as a free parameter to a model comparison using the cbm toolkit. This comparison showed that the original fmUBC, absent any mechanism to carry learned values across contexts, fit the data best (exceedance probability = 0.97), showing that participants did indeed adhere to the instructed task structure.

This comparison across computational models offers evidence that participants did not carry values across contexts as they were instructed to do. However, we wanted to ensure that the influence of previous context wasn’t simply misattriuted and accommodated by other mechanisms in the model. We also wanted to probe for timepoints within a block of trials where choice might exhibit the influence from previous contexts. To address these concerns we implemented a sliding window regression analysis, conducting a model comparison between a model that included values from previously encountered contexts to a model that didn’t. We defined a baseline model as:

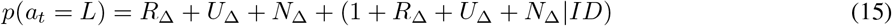

where *R*_∆_ = *E*[*S*_*L*_] − *E*[*S*_*R*_] denotes the reward differential in favor of the left option, while *U*_∆_ = *U* [*S*_*L*_] − *U* [*S*_*R*_], and *N*_∆_ = *N* [*S*_*L*_] − *N* [*S*_*R*_] reflect the difference in uncertainty and novelty respectively, with random intercept and slopes estimated for each participant ID, with values defined as they were for Equation 1. We then defined an augmented model that also included the expected value leaned from the most recent context:

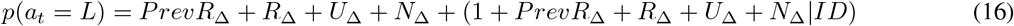

where *PrevR*_∆_ denotes the previous context’s expected value differential between left and right stimuli. We then repeatedly fit and compared the variance explained by both model across a sliding window of two trials within blocks (e.g. window 1 = trials 1 and 2, window 2 = trials 2 and 3, etc…). This analysis showed that the previous value did not offer sufficient additional explanatory power for any window of trials (all p-values > 0.05, uncorrected for multiple comparisons).

In summary, neither computational model comparison, nor computationally agnostic analysis of fine grained structure embedded in participant’s game-play showed evidence of value carry-over from previous contexts, demonstrating that participant behavior reflected the instructed task structure faithfully.

### 5.2 Other experimental variables not predictive of novelty seeking

Participants exhibited a robust preference for novel options that increased as they progressed through a learning context. The fmUCB model reproduces this phenomena by combining a constant novelty bias pulling participants toward novel options, and a growing push away from paired uncertain familiar options. However, colloquial interpretations such as growing boredom, or a superstition that that novel options might offer a bonus may also tempt inquiry.

Participants may have adopted the unfounded belief that novel options were baited to offer a bonus. Participants were not offered any instructions to hint at this, nor was there an empirical difference in the rewards experienced after sampling a novel or familiar option for the first time (mean reward for both familiar and novel options = 0.5, *t*(31) = −0.2, *p* = 0.85). Furthermore, given that participants exhibited a growing preference for novel options, they would presumably need to assume that novel options presented later in a block of trials was more likely to offer a baited bonus than novel options presented earlier. No participants reported such a strategy or belief.

Participants performed the experimental task for approximately 75 minutes while in the scanner, with an average time of just under 4 minutes per learning block. Most participants reported finding the task challenging and relatively fun. However, to demonstrate that behavior wasn’t influence by fatigue, boredom, or other anomalous time-on-task phenomena, we probed for the emergence of shifting strategies as the experimental task progressed. We modified the regression model defined in Equation 1 to include block number as an interaction term instead of trial number.

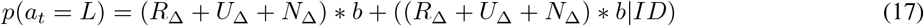

Model comparison showed that trial number offered a significantly better predictor of choice than block number (log-likelihoods -6515.5 and -6557.9 respectively), demonstrating that preferences for novelty, uncertainty or rewarded stimuli did not shift meaningfully as the experiment progressed. We constructed a second comparative model by augmenting the model described by Equation 1 to include an additional term specifically probing for an effect of block on novelty seeking:

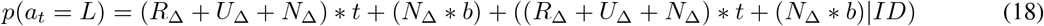

Replicating effects previously reported, this model identified a significant novelty seeking bias (*β*_*N*_ = 0.18, *p* < 0.01) that increased within the block of trials (*β*_*N*:*t*_ = 0.42, *p* < 0.01). However, there was no effect of block number (*β*_*b*_ =− 0.08, *p* = 0.44) nor was there a significant interaction with novelty (*β*_*N*:*b*_ = −0.05, *p* = 0.6). Furthermore, model comparison showed that the additional block number variable was unwarranted (*χ*^2^ (21, 32) = 20.528, *p* = 0.49), demonstrating that novelty seeking strategies are not accounted for by experiment duration (as opposed to block level) variables.

**Supplementary Figure 1:**
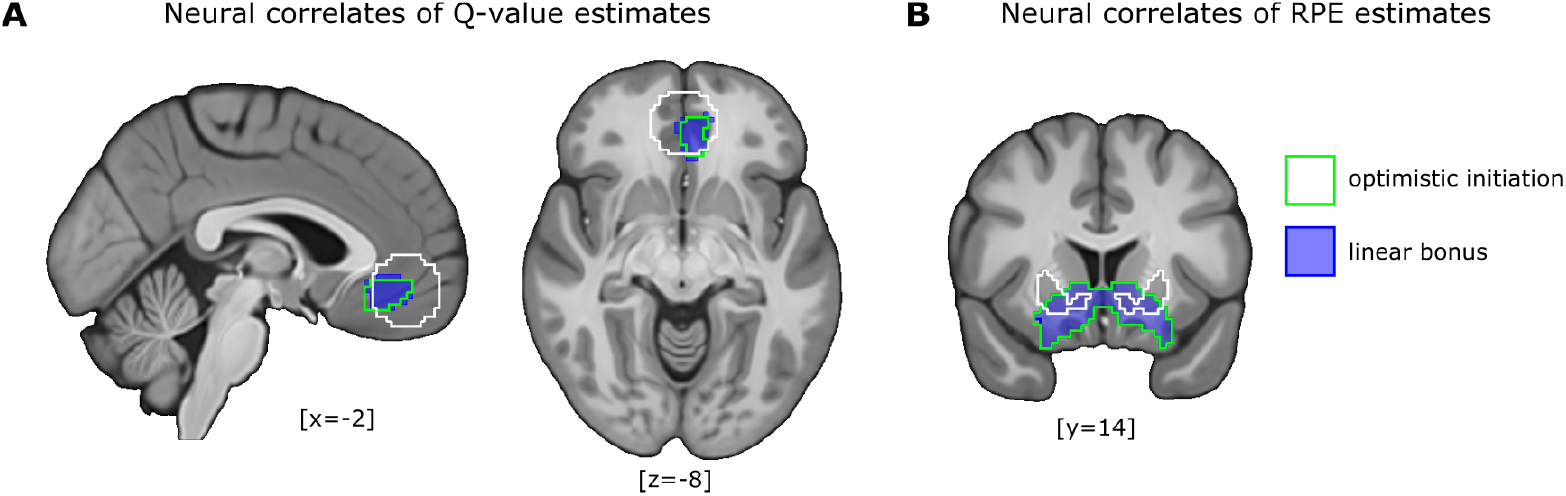
Voxels correlated with expected value and reward prediction errors across mechanisms of novelty integration. **A)** A cluster of voxels in vmPFC correlated with q-values estimated by the linear novelty bonus mechanism (blue), and voxels correlated with q-values as estimated by the optimistic initiation mechanism (green bordered). Mask comprising 15mm sphere centered on independent meta-analysis peak voxel in vmPFC bordered in white. **B)** A cluster of voxels in ventral striatum correlated with reward prediction errors estimated by the linear novelty bonus mechanism (blue), and voxels correlated with reward prediction errors as estimated by the optimistic initiation mechanism (green bordered). Whole-brain maps for signal of interest were tested with a cluster-forming threshold of *P* < 0.001 uncorrected, followed by cluster-level FWE correction at P<0.05.

### 5.3 Behavioral confusability analysis

Discriminability between the optimistic initiation and the linear bonus mechanism was probed via model confusability. We first fit the fmUCB model defined in terms of both the optimistic initiation and the linear bonus mechanisms to participant data, and used those optimized parameter estimates to specify each model’s free parameters. Each model instantiation was then exposed to the same set of trials experienced by our participants, and generated a choice on each trial. This process was repeated 100 times for each participant, resulting in 100 simulated experiments with data generated by each of the two mechanisms. Finally, we fit both implementations to both set of simulated experiments to quantify the proportion of fits that correctly identify the true generative mechanism. This analysis revealed that data generated by the optimistic initiation mechanism was only correctly identified for 53% ± 0.14 of the fits. Data generated by the linear bonus mechanism suffered equally poor identification, with 57 % ± 1.8 of the fits correctly identifying the generative mechanism. Thus, we conclude that behavior alone cannot distinguish between either the optimistic or linear novelty bonus mechanisms given our experimental design.

### 5.4 Neural correlates of expected reward associated with the linear bonus model

We derive estimated time-courses for variables of interest by fitting the fmUCB model defined to use the linear novelty bonus mechanisms (see Equation 12) to participant behavior. We then applied model estimates as parametric regressors in the GLM outlined in Section 2.6 to identify regions associated with stimulus reward valuation (selected + rejected q-value). This analysis identified a cluster in vmPFC that encompassed the cluster associated with q-values estimated by the fmUCB model (see Figure 1A). Regions identified by both models are subsumed by the vmPFC ROI independently identified by meta-analyses [27, 28]. An analysis of regions correlated with the reward prediction errors generated by both models also revealed largely overlapping correlates in ventral striatum (see Figure 1B).

### 5.5 Neural correlates of the uncertainty bonus absent familiarity modulation

The GLM described in Section 2.6 was modified to include the uncertainty bonus term estimated by the fmUCB model absent familiarity modulation to identify regions associated with value (selected + rejected uncertainty bonus):

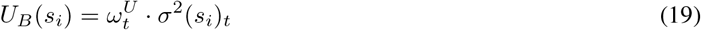

No significant clusters were found using conventional voxel height p<0.001, but we report a cluster at an extremely liberal threshold of p<0.01. Notably, this cluster overlaps with the region associated with the familiarity modulated uncertainty bonus reported in Figure 4B.

**Supplementary Figure 2:**
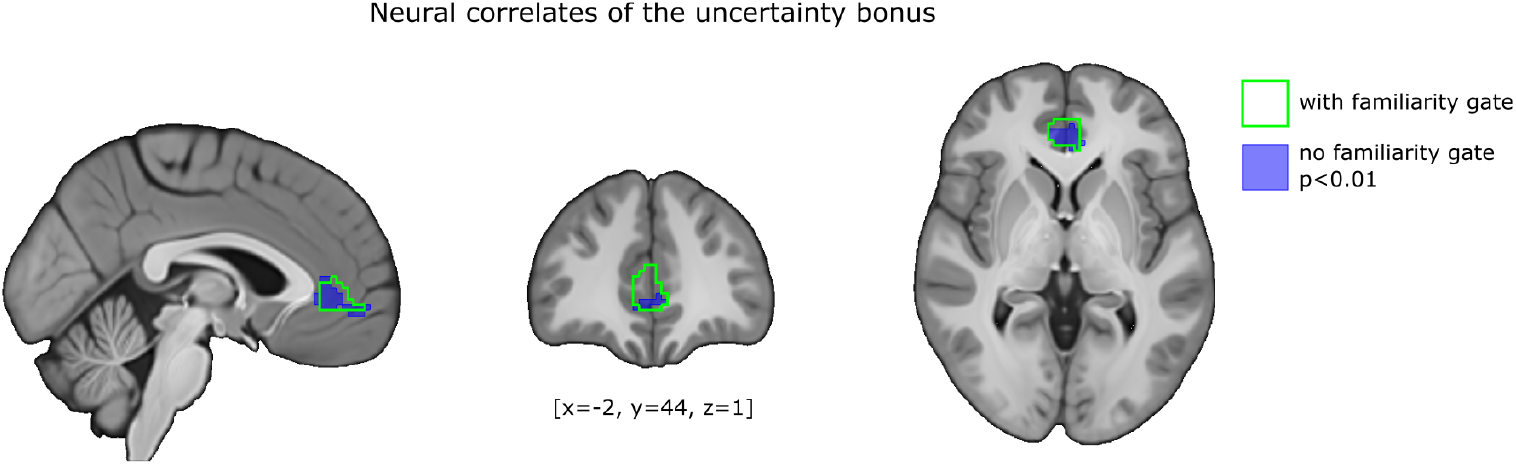
Voxels correlated with the uncertainty bonus. A cluster of voxels in mPFC correlated with the uncertainty bonus absent familiarity modulation (blue), but only at an extremely liberal threshold of p<0.01. This region overlaps the cluster associated with the familiarity modulated uncertainty bonus (green border), which was tested with a cluster-forming threshold of *P* < 0:001 uncorrected, followed by cluster-level FWE correction at *P*<0.05.

## 6 Acknowledgements

This work was funded by grants from the US National Science Foundation (1207573) to JOD, the Natural Science and Engineering Research Council (RGPIN-2018-05946) to WC, and the Social Sciences and Humanities Research Council (CGS-MSFSS) to VM.

## 7 Author information

### 7.2 Contributions

J.C and J.P.O conceived of the experiments and data analysis. J.C, J.P.O, V.M, and W.C wrote the manuscript. J.C carried out the fMRI experiment, and V.M carried out the replication study.

## References

[1] Jonathan D Cohen, Samuel M McClure, and Angela J Yu. Should i stay or should i go? how the human brain manages the trade-off between exploitation and exploration. Philosophical Transactions of the Royal Society B: Biological Sciences, 362(1481):933–942, 2007.

[2] John C Gittins. Bandit processes and dynamic allocation indices. Journal of the Royal Statistical Society: Series B (Methodological), 41(2):148–164, 1979.

[3] Abdelkader Ennaceur and Jean Delacour. A new one-trial test for neurobiological studies of memory in rats. 1: Behavioral data. Behavioural brain research, 31(1):47–59, 1988.

[4] Robert N Hughes. Neotic preferences in laboratory rodents: issues, assessment and substrates. Neuroscience & Biobehavioral Reviews, 31(3):441–464, 2007.

[5] Kirk R Daffner, M Marsel Mesulam, Leonard FM Scinto, Lisa G Cohen, Bruce P Kennedy, W Caroline West, and Phillip J Holcomb. Regulation of attention to novel stimuli by frontal lobes: an event-related potential study. Neuroreport, 9(5):787–791, 1998.

[6] Bianca C Wittmann, Nathaniel D Daw, Ben Seymour, and Raymond J Dolan. Striatal activity underlies novelty-based choice in humans. Neuron, 58(6):967–973, 2008.

[7] Robert C Wilson, Andra Geana, John M White, Elliot A Ludvig, and Jonathan D Cohen. Humans use directed and random exploration to solve the explore–exploit dilemma. Journal of Experimental Psychology: General, 143(6):2074, 2014.

[8] David Badre, Bradley B Doll, Nicole M Long, and Michael J Frank. Rostrolateral prefrontal cortex and individual differences in uncertainty-driven exploration. Neuron, 73(3):595–607, 2012.

[9] Samuel J Gershman. Deconstructing the human algorithms for exploration. Cognition, 173:34–42, 2018.

[10] Tommy C Blanchard and Samuel J Gershman. Pure correlates of exploration and exploitation in the human brain. Cognitive, Affective, & Behavioral Neuroscience, 18(1):117–126, 2018.

[11] Nadescha Trudel, Jacqueline Scholl, Miriam C Klein-Flügge, Elsa Fouragnan, Lev Tankelevitch, Marco K Wittmann, and Matthew FS Rushworth. Polarity of uncertainty representation during exploration and exploitation in ventromedial prefrontal cortex. Nature Human Behaviour, pages 1–16, 2020.

[12] Rajeev Agrawal. Sample mean based index policies with o (log n) regret for the multi-armed bandit problem. Advances in Applied Probability, pages 1054–1078, 1995.

[13] Michael N Katehakis and Herbert Robbins. Sequential choice from several populations. Proceedings of the National Academy of Sciences of the United States of America, 92(19):8584, 1995.

[14] Peter Auer, Nicolo Cesa-Bianchi, and Paul Fischer. Finite-time analysis of the multiarmed bandit problem. Machine learning, 47(2-3):235–256, 2002.

[15] Ronen I Brafman and Moshe Tennenholtz. R-max-a general polynomial time algorithm for near-optimal reinforcement learning. Journal of Machine Learning Research, 3(Oct):213–231, 2002.

[16] Andrew Y Ng, Daishi Harada, and Stuart Russell. Policy invariance under reward transformations: Theory and application to reward shaping. In ICML, volume 99, pages 278–287, 1999.

[17] Anne GE Collins and Jeffrey Cockburn. Beyond dichotomies in reinforcement learning. Nature Reviews Neuroscience, 21(10):576–586, 2020.

[18] Ifat Levy, Jason Snell, Amy J Nelson, Aldo Rustichini, and Paul W Glimcher. Neural representation of subjective value under risk and ambiguity. Journal of neurophysiology, 103(2):1036–1047, 2010.

[19] Elise Payzan-LeNestour, Simon Dunne, Peter Bossaerts, and John P O’Doherty. The neural representation of unexpected uncertainty during value-based decision making. Neuron, 79(1):191–201, 2013.

[20] Nathaniel D Daw, John P O’doherty, Peter Dayan, Ben Seymour, and Raymond J Dolan. Cortical substrates for exploratory decisions in humans. Nature, 441(7095):876–879, 2006.

[21] Michael J Frank, Bradley B Doll, Jen Oas-Terpstra, and Francisco Moreno. Prefrontal and striatal dopaminergic genes predict individual differences in exploration and exploitation. Nature neuroscience, 12(8):1062, 2009.

[22] Jan-Benedict EM Steenkamp and Katrijn Gielens. Consumer and market drivers of the trial probability of new consumer packaged goods. Journal of Consumer Research, 30(3):368–384, 2003.

[23] Wako Yoshida and Shin Ishii. Resolution of uncertainty in prefrontal cortex. Neuron, 50(5):781–789, 2006.

[24] Erie D Boorman, Timothy EJ Behrens, Mark W Woolrich, and Matthew FS Rushworth. How green is the grass on the other side? frontopolar cortex and the evidence in favor of alternative courses of action. Neuron, 62(5):733–743, 2009.

[25] Silvia A Bunge and Carter Wendelken. Comparing the bird in the hand with the ones in the bush. Neuron, 62(5):609–611, 2009.

[26] Anjali Raja Beharelle, Rafael Polanía, Todd A Hare, and Christian C Ruff. Transcranial stimulation over frontopolar cortex elucidates the choice attributes and neural mechanisms used to resolve exploration–exploitation trade-offs. Journal of Neuroscience, 35(43):14544–14556, 2015.

[27] John A Clithero and Antonio Rangel. Informatic parcellation of the network involved in the computation of subjective value. Social cognitive and affective neuroscience, 9(9):1289–1302, 2014.

[28] Oscar Bartra, Joseph T McGuire, and Joseph W Kable. The valuation system: a coordinate-based meta-analysis of bold fmri experiments examining neural correlates of subjective value. Neuroimage, 76:412–427, 2013.

[29] Philippe Domenech, Sylvain Rheims, and Etienne Koechlin. Neural mechanisms resolving exploitation-exploration dilemmas in the medial prefrontal cortex. Science, 369(6507), 2020.

[30] Jon C Horvitz, Tripp Stewart, and Barry L Jacobs. Burst activity of ventral tegmental dopamine neurons is elicited by sensory stimuli in the awake cat. Brain research, 759(2):251–258, 1997.

[31] P Read Montague, Peter Dayan, and Terrence J Sejnowski. A framework for mesencephalic dopamine systems based on predictive hebbian learning. Journal of neuroscience, 16(5):1936–1947, 1996.

[32] Wolfram Schultz. Predictive reward signal of dopamine neurons. Journal of neurophysiology, 80(1):1–27, 1998.

[33] Sham Kakade and Peter Dayan. Dopamine: generalization and bonuses. Neural Networks, 15(4-6):549–559, 2002.

[34] Ruth M Krebs, Björn H Schott, Hartmut Schütze, and Emrah Düzel. The novelty exploration bonus and its attentional modulation. Neuropsychologia, 47(11):2272–2281, 2009.

[35] Payam Piray, Amir Dezfouli, Tom Heskes, Michael J Frank, and Nathaniel D Daw. Hierarchical bayesian inference for concurrent model fitting and comparison for group studies. PLoS computational biology, 15(6):e1007043, 2019.

[36] Lotem Elber-Dorozko and Yonatan Loewenstein. Striatal action-value neurons reconsidered. ELife, 7:e34248, 2018.

[37] John P O’Doherty, Peter Dayan, Karl Friston, Hugo Critchley, and Raymond J Dolan. Temporal difference models and reward-related learning in the human brain. Neuron, 38(2):329–337, 2003.

[38] Wojciech K Zajkowski, Malgorzata Kossut, and Robert C Wilson. A causal role for right frontopolar cortex in directed, but not random, exploration. Elife, 6:e27430, 2017.

[39] Thomas J Palmeri, Bradley C Love, and Brandon M Turner. Model-based cognitive neuroscience. Journal of Mathematical Psychology, 76:59–64, 2017.

[40] Richard Henson. What can functional neuroimaging tell the experimental psychologist? The Quarterly Journal of Experimental Psychology Section A, 58(2):193–233, 2005.

[41] Mike PA Page. What can’t functional neuroimaging tell the cognitive psychologist? Cortex, 42(3):428–443, 2006.

[42] Todd A Hare, Colin F Camerer, and Antonio Rangel. Self-control in decision-making involves modulation of the vmpfc valuation system. Science, 324(5927):646–648, 2009.

[43] Shinsuke Suzuki, Logan Cross, and John P O’Doherty. Elucidating the underlying components of food valuation in the human orbitofrontal cortex. Nature neuroscience, 20(12):1780–1786, 2017.

[44] John P O’Doherty, Jeffrey Cockburn, and Wolfgang M Pauli. Learning, reward, and decision making. Annual review of psychology, 68:73–100, 2017.

[45] Stephen M Smith, Mark Jenkinson, Mark W Woolrich, Christian F Beckmann, Timothy EJ Behrens, Heidi Johansen-Berg, Peter R Bannister, Marilena De Luca, Ivana Drobnjak, David E Flitney, et al. Advances in functional and structural mr image analysis and implementation as fsl. Neuroimage, 23:S208–S219, 2004.

[46] Brian B Avants, Nick Tustison, and Gang Song. Advanced normalization tools (ants). Insight j, 2(365):1–35, 2009.

[47] J Michael Tyszka and Wolfgang M Pauli. In vivo delineation of subdivisions of the human amygdaloid complex in a high-resolution group template. Human brain mapping, 37(11):3979–3998, 2016.

[48] William D Penny, Karl J Friston, John T Ashburner, Stefan J Kiebel, and Thomas E Nichols. Statistical parametric mapping: the analysis of functional brain images. Elsevier, 2011.

